# Honeybee cognition as a tool for scientific engagement

**DOI:** 10.1101/2021.05.08.442068

**Authors:** Jai A. Denton, Ivan Koludarov, Michele Thompson, Jarosław Bryk, Mariana Velasque

**Affiliations:** Genomics & Regulatory Systems Unit, Okinawa Institute of Science & Technology, Tancha, Okinawa, Japan; Institute of Vector-borne Disease, Monash University, Clayton, Victoria, Australia; Animal Venomics Group, Justus Liebig University, Giessen, Hessen, Germany; Independent Researcher, Dexter, Michigan USA; School of Applied Sciences, University of Huddersfield, Huddersfield, UK

**Keywords:** citizen science, honeybee learning, memory, proboscis extension response, associative learning, honeybee cognition

## Abstract

In addition to the crowdsourcing of experimental data, citizen science, and scientific engagement more broadly, serve as a bridge between researchers and the wider community. This serves to foster a greater understanding of the scientific method and science-based solutions generally. *Apis mellifera* (honeybees) are a well-established model for the study of learning and cognition and can serve as an engaging outreach system for this wider community. Here, we developed and implemented a protocol using well established honeybee conditioning protocols to safely study the effects of caffeine and dopamine on learning performance. Using this protocol, a group of high-school aged students as part of the Ryukyu Girls program demonstrated that caffeine, but not dopamine, significantly reduced the number of trials required for a successful conditioning response. This allowed these students to explore the scientific method in a relatable and engaging way.

**Simple Summary:** Global scientific literacy can be improved through widespread and effective community engagement by researchers. We propose *Apis mellifera* (honeybee) as an public engagement tool due to widespread awareness of colony collapse and the bees’ importance in food production. Moreover, their cognitive abilities make for engaging experiments. Their relative ease of cultivation means that studies can be performed cost-effectively, especially when partnering with local aperists. Using a proxy for honeybee learning, a group of non-specialist high-school-aged participants obtained data suggesting that caffeine, but not dopamine, improved learning. This hands-on experience facilitated student understanding of the scientific method, factors that shape learning and the importance of learning for hive health.

## 1. Introduction

We currently face an unprecedented combination of events from pandemics and climate emergency to desertification of arable lands and the extinction of predators, the challenges whose understanding and overcoming demands broad scientific expertise. Thus, it is imperative that communities not only understand the scientific method, but also engage with it [7–9]. A key approach to drive this engagement is through citizen science activities. Although there is not a single concise definition of citizen science, it can typically be thought of as involving non-scientists in the scientific process. However, many of the most significant programs either engage with community members already scientifically literate or are much more goal-focused without necessarily improving the understanding of the scientific method [10–12]. Therefore, effective community engagement and citizen science needs to include both tentpole and smaller activities with community members that are underrepresented in the scientific process.

Although citizen science and scientific outreach take many forms, it is most effective when participants can meaningfully engage with the topic [10,13,14]. The central activity, experiment or scientific question needs to incite enthusiasm but also needs to be presented in a framework that ensures participants gain the most from the experience [10,13,14]. Ultimately, citizen science programs are not only beneficial for students, but also help to build positive relationships between universities that sponsor them and the local community. Due to the broad appeal, external funding from private donors and corporate grants, with more flexible spending parameters, are often available and can be easier to secure.

The Ryukyu Girls scientific engagement program seeks to engage female high school students from the Japanese Okinawa prefecture in the scientific process. As part of this program, we developed an experiment that sought to teach the scientific method as a way of interacting with the world. We employed clear and simple language when describing the scientific processes to ensure it was accessible to non-native English speakers. The program was taught in two languages, English and Japanese, with simultaneous translation provided. We have also gone beyond the language when our presenter made purposeful choices to forge a human connection by sharing their life experiences. As a result, the participants were not only more deeply engaged with the content, but also developed a level of trust with the presenter, who they clearly viewed as a role model.

In addition, a critical component of our approach was in highlighting the importance of participation in the scientific process. By drawing a connection between their participation as a part of a larger effort to increase underrepresented voices in science, participants became emotionally invested and, as a result, experienced meaningful engagement in the experiment.

*Apis mellifera* (European honeybees) provides an excellent tool for scientific engagement, of academic and lay public alike, while furthering our understanding of this critical agricultural pollinator. Honeybees have complex social interactions driven by intra-hive learning and communication [15]. They are capable of not only learning the location of food resources but also communicating to their nestmates through waggle dance [15,16]. In the laboratory, honeybees are exceptional model organisms to study cognition, memory and communication. Honeybees are capable of complex cognitive processes. For instance, honeybees can memorise locations, patterns, faces and even understand conceptual relationships, such as above/below and same/different [17,18]. Despite this great potential for honeybees as models in cognitive neuroscience, its use is still limited when compared to *Drosophila melanogaster* [19].

Classical conditioning is a form of conditioning on which a subject learns to associate a neutral stimulus, called conditioned stimulus (CS), with a stimulus of biological significance, the unconditioned stimulus (US), such as sucrose [20]. Over time, animals start to associate the initial neutral stimulus to the US, acquiring the capacity to elicit a conditioned response. Although classical conditioning is considered to be a basic learning process, it has become the foundation of cognition and memory studies in animals, especially in insects [21–26]. Amongst insects, *Apis mellifera* is considered one of the most robust organisms for the study of classical conditioning [27–30]. Such success is mainly due to the presence of several powerful conditioning protocols [24,25,29,31–34].

Honeybees extend their proboscis when their chemoreceptors enter in contact with sucrose. When sucrose is paired with another stimulus, such as a distinctive scent or a visual pattern, honeybees can learn to anticipate the sugar reward when exposed to the stimulus, extending their proboscis. In conditioning, a naive bee (i.e. a honeybee without any previous experience to the stimulus) is exposed to a neutral stimulus (i.e. a scent or an image it has not being exposed to before), CS, followed by a sucrose reward, US [24,25,29,31,32]. During conditioning, the honeybee learns to associate the initially neutral stimulus, CS, to the US (i.e. sucrose). Although honeybees are amenable to cognitive study, cost-effective, and ubiquitous worldwide, conditioning protocols rely on the implementation of several procedural steps [2]. Such steps, such as training trials, can be laborious and require large sample sizes, which can be an impediment to obtaining statistically robust findings [35]. Therefore, the presence of a system that provides large-scale data in a semi-structured system, through citizen science activities and/or through automation, can help to overcome these challenges.

Herein we describe the use of *A. mellifera* learning as an experimental system by investigating how dopamine and caffeine affects learning performance in honeybees. The study was implemented by Ryukyu Girls engagement program, as an example of outreach and citizen science activities. The results indicated that caffeine-treated bees learn faster than dopamine-treated and control bees. All data was collected and analysed by the participants of the programme, with little data loss and with results comparable to the pilot trial, suggesting that the protocol and the programme such as Ryukyu Girls in particular can be used to support large-scale observation and data collection.

## 2. Methods & Protocol

### 2.1 Honeybee Sourcing and Handling

All bees used were from a single hive. Newly emerged bees were obtained by removing two frames containing capped larvae from the hive, brushing off adult bees and placing the frames in a small hive box for 6 hours. To maintain optimal conditions for honeybees, small hives were kept inside the incubator, with constant temperature of 34°C and 60% humidity. After 6 hours all emerged honeybees were collected and harnessed for experimentation. The two frames were returned to the hive as soon as possible after removing the newly emerged bees.

### 2.2 Harnessing

After being anesthetized using ice [36], bees were harnessed using plastic drinking straws, approximately 14mm in diameter, and cut to lengths of approximately 2cm with a diagonal section removed from one end, creating a V, to allow the bee’s head to protrude as described by [37]. Anesthetized bees were placed inside the straw piece, restricting the movement of their legs. The head was fixed in place with small pieces of masking tape that allowed movement of the antennae and proboscis. Although this process is well described by [2] et al. [2], harnessing requires patience and practice to effectively perform. Harnessed bees were divided in three groups, providing three bees per treatment group to the students. To facilitate handling and reduce confusion associated with the treatment, harnessed bees were placed in small holes present in a styrofoam tray, containing a color coded indication and names of the treatment groups (Figure 1a). To reduce mortality and stress, bees were harnessed 6 hours prior to the practical by the authors without the support of the Ryukyu Girls class.

**Figure 1.**
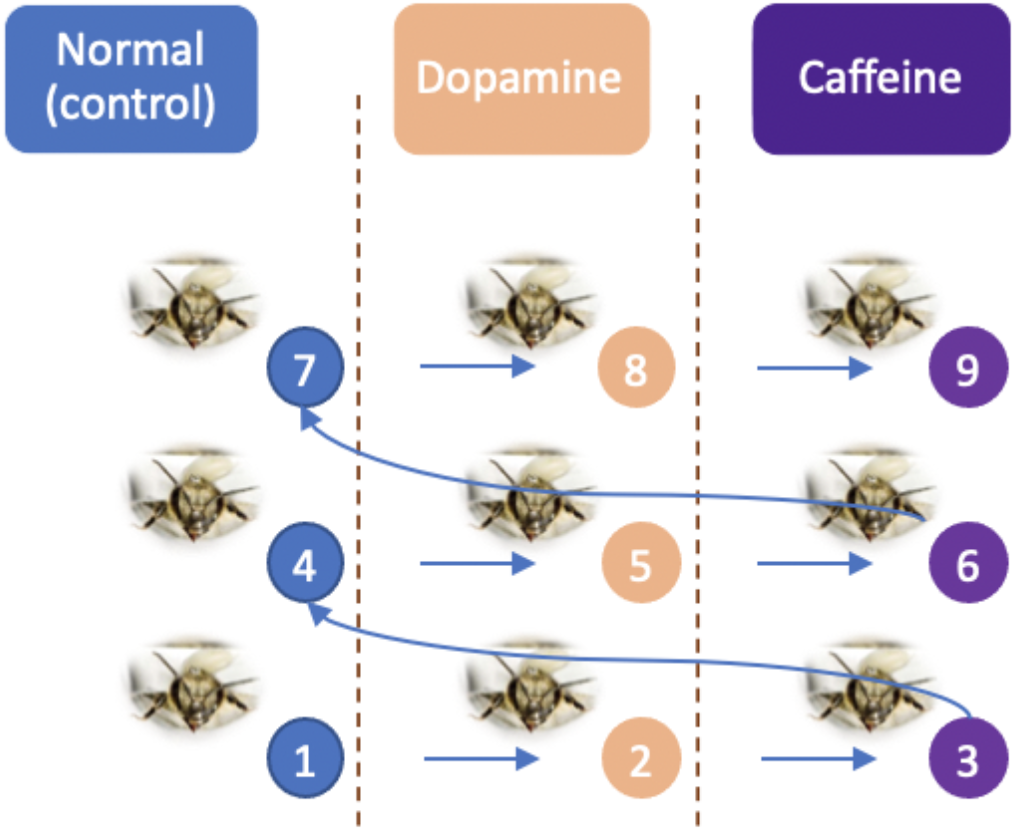
Diagram of experimental design and workflow provided in the laboratory notebook. The workflow delineates the direction of movement, with the first individuals of each treatment group completing the training trial before proceeding to the second and so on, returning to the first individual upon completion.

### 2.3 Quantifying Learning

Despite honeybees being a well established model organism, there is mixed evidence to whether newly emerged bees are able to show learning [1]. Therefore, a pilot experiment was conducted to ensure newly emerged honeybees would be capable of associative learning.

Learning performance was measured as the number of trials required for the honeybee to learn a new stimulus. In this case, bees were exposed to a lemon scent while offering the sugar solution. When offered a sugar syrup reward, bees extend their proboscis. Over a few trials (i.e. odour paired with sugar reward) the bee learns to associate the odour to the reward, extending their proboscis when exposed to the odor alone. Fewer trials indicates higher learning performance.

### 2.4 Pilot Training

Pilot was performed in 18 bees (6 bees per treatment group) and allowed to establish the presence of associative learning in newly emerged honeybees and if there was a clear trend according to the treatment group.

### 2.5 Pre-Experimental Preparation

Because dopamine and caffeine can vary on the time to have an effect, depending on the organism and age, all bees were fed three times prior to the experiment. Feeding was done at 2, 1 and 0.5 hours before the experiment with a sucrose solution mixed with the proposed treatment. Bees were fed according to their experimental group, with sucrose (sucrose solution only: control) or sucrose mixed with either caffeine (sucrose + caffeine: caffeine) or dopamine (sucrose + dopamine: dopamine) solution. The feeding was not paired with odour cue.

Sucrose solution was prepared by mixing 200ml water with 100g of sucrose and stirred until all sugar was dissolved. The sucrose solution was divided in three equal parts. One part was mixed with 1mg of caffeine (caffeine treatment; ALLMAX Nutrition) and another with 1mg of dopamine (dopamine treatment; Dopamine Plus N-care). Both solutions were stirred until the chemical was dissolved. All solutions were allocated in 1ml eppendorf tubes to be used during the practice.

To create the odour stimulus during the trials, one drop of lemon essential oil (Lemon Essential Oil; My Pure Earth) in a cotton ball was used. To prevent evaporation and further contamination, the cotton ball was stored inside an eppendorf tube. All solutions used during the pre-preparation stage and during experimental training were prepared 18h prior to the practice. All material used in the experimental training was labelled and colour coded, following the same styrofoam tray scheme (Figure 1).

### 2.6 Experimental Training

Prior to the experiment the Ryukyu Girls Class were provided with an explanation of the motivation, literature review (i.e. effects of dopamine in humans and other animals) and what a hypothesis is. Following, they were shown the experimental setup and divided in groups of three. Each group received 3 copies of the “laboratory notebook” (see supplemental material for a copy of the laboratory notebook), eppendorf tubes containing the treatment solutions, the odour scent and multiple swabs. They were then asked to formulate a hypothesis and a prediction for the experiment. They were informed about the risks related to honeybees and how to proceed in case they escape their harness.

The training trial consisted of approximating the eppendorf tube containing the lemon scent near the honeybee antennae followed by lightly touching the antennae with the sucrose cotton swab until the bee extends its proboscis, followed by allowing them to feed for one second (Figure 2). The sucrose cotton swab contained either dopamine, caffeine or just sucrose solution, according to the bee experimental group. Each trial was then repeated across all groups following the diagram present in the Figure 1b. They were instructed to repeat the trial multiple times. At the third training trial, students were instructed to delay the sucrose reward for a few seconds to allow visualization of the associative learning. Such small delay, would allow students to visualise proboscis extension without impairing further training protocol. Therefore, learning would be accounted for only after the third training trial.

**Figure 2.**
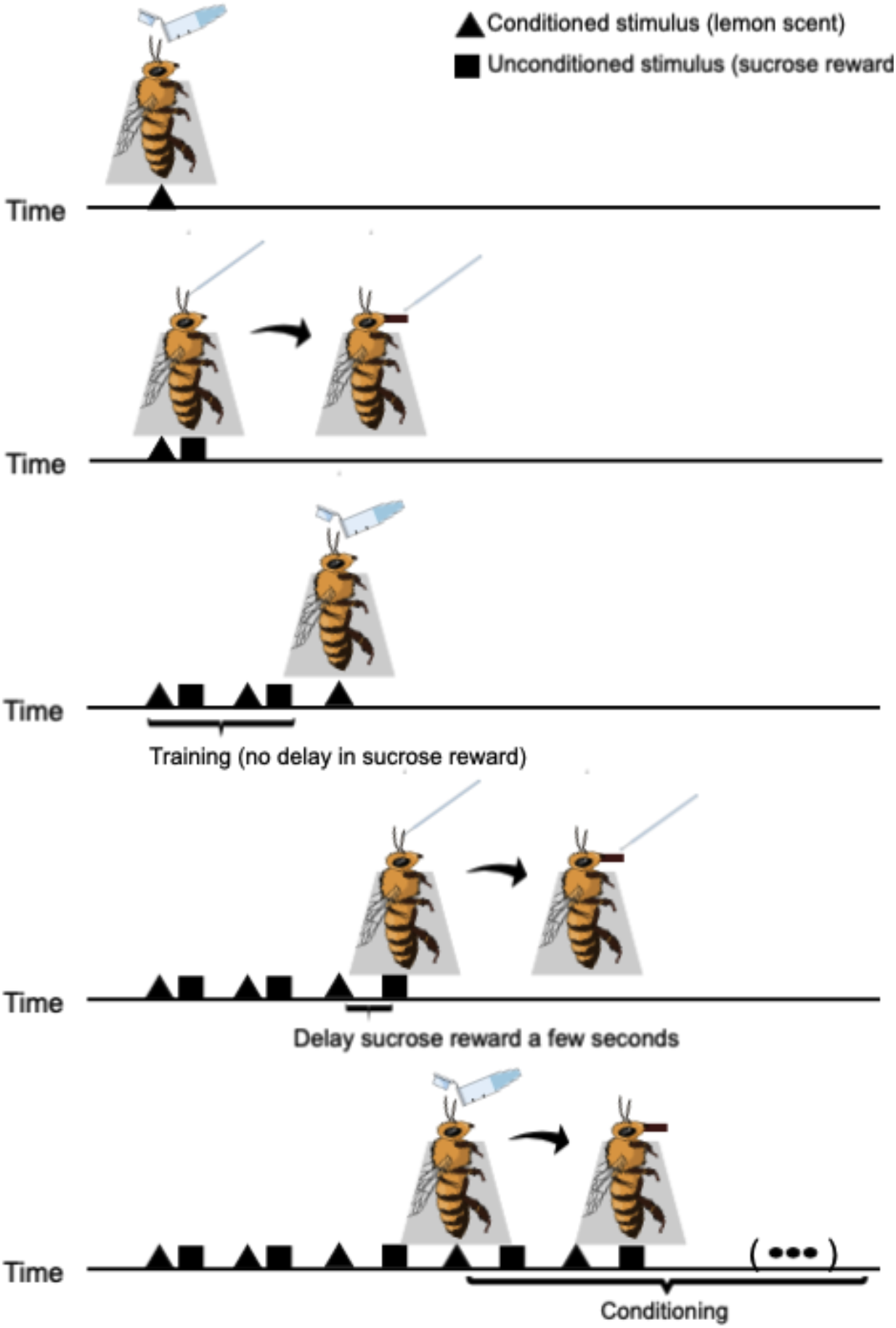
The training schematic for the conditioning of honeybees using a lemon scent. The number of conditioned/unconditioned stimuli cycles can be varied. The conditioning was repeated across all treatments

We considered that the bee learned when they exposed their proboscis to the lemon scent prior to the offer of the sucrose solution. The number of trials per bee per group was recorded in the lab notebook and supplied to the authors for posterior analysis.

### 2.7 Cost

Around 100 dollars for the hive supplied. The use of newly emerged bees for the practice did not cause significant damage to the hive and it could be used in further experiments.

### 2.8 Student Group and Engagement

To facilitate learning and increase student engagement, we developed a powerpoint presentation and a lab notebook. Both the presentation and lab notebook contained the research background and the protocol for the experiment (lab notebook and presentation can be found on the supplemental materials).

### 2.9 Risk Management

When employing an experimental system that poses a stinging and subsequent allergy risk diligence is required. We employed several strategies to ensure no participants were stung. Although we conducted an initial survey of participants regarding their allergy status to honeybees, this is insufficient to minimise risk. Where possible, double containment was employed. We used only european newly emerged bees, less than 20 hours old, as they are both less aggressive and do not produce venom [38]. We also developed a harness-like system (see Harnessing) that prevents contact with the bee abdomen. We ensured staff levels were such that every experimental group could be supervised by a demonstrator so if a bee did escape, it could be quickly caught. As a final precaution we also kept an adrenaline injector (EpiPen) in our medical kit.

### 2.10 Data analysis

Behaviour is one of the most labile phenotypes and thus use of an appropriate statistical analysis is required to accommodate experimental variation. To minimise issues related to handling, repeated measures ANOVA was employed to analyse the effect of caffeine and dopamine on learning performance. Repeated measures ANOVA compares the difference between means across the treatment groups that are based on repeated observations. Differences between groups were estimated using Multiple Comparisons of Means (Tukey Contrasts). All data analysis was performed in R (R version 4.0.2) and can be viewed at: https://github.com/marivelasque/HoneybeeOutreach.git

## 3. Results

### 3.1 Hypothesis Development

To facilitate engagement and understanding, considerable classroom time focused on the process of developing a hypothesis and subsequent testing. Although due to time and logistical constraints, hypotheses could not be generated spontaneously within the classroom. However, after describing the context, materials available and providing guidance in the form of classroom discussion, participants developed hypotheses nearly identical to those we sought to test.

### 3.2 Data Reliability, Collation and Analysis

A total of twelve student groups performed each experiment in triplicate. However, due to loss of bees or experimental error, the groups averaged 2.4 bees per treatment. Each experimental treatment comprises, on average, 28.6 data points. Of the total 108 data points collected, 22 failed. This data loss was predominantly related to harnessing of the bees (see Harnessing). Newly emerged bees have softer cuticles than adults [39] and thus, improper handling during harnessing and training could be partially responsible for the high mortality.

Subsequent to completing the experimental work, data collation and a brief analysis was conducted within a classroom setting. This was closely linked to the aforementioned hypothesis generation and thus provided participants with a further understanding in hypothesis testing.

### 3.3 Group results

Caffeine, but not dopamine, was found to significantly reduce the number of trials required for a successful conditioning response (Table 1; Figure 3; Supplementary Table 1). Each of the 12 groups conducted the control, caffeine treatment and dopamine treatment, in triplicate, by measuring the number of trials required for a conditioning response. Although the average was lower for both caffeine and dopamine treatments, only caffeine had a statistically significant difference (caffeine p = 0.038, dopamine p = 0.252; repeated measures anova).

**Table 1.** Collated data from the 12 student groups. Values indicate the average number of trials required for conditioning response.

| Group ID | Control | Dopamine | Caffeine |
| --- | --- | --- | --- |
| 1 | 6.00 | 5.67 | 4.50 |
| 2 | 4.00 | 5.00 | 5.67 |
| 3 | 6.33 | 6.00 | 4.00 |
| 4 | 5.00 | 7.00 | 5.00 |
| 5 | 5.00 | 4.50 | 5.00 |
| 6 | 4.00 | 4.00 | 4.50 |
| 7 | 5.00 | 6.50 | 5.00 |
| 8 | 6.33 | 7.00 | 4.00 |
| 9 | 4.67 | 5.00 | 5.67 |
| 10 | 6.00 | 6.00 | 6.00 |
| 11 | 5.00 | 6.67 | 5.00 |
| 12 | 4.00 | 6.33 | 4.67 |
| **Average** | 5.11 | 5.81 | 4.92 |
| **St. Dev.** | 0.84 | 0.94 | 0.61 |

**Figure 3.**
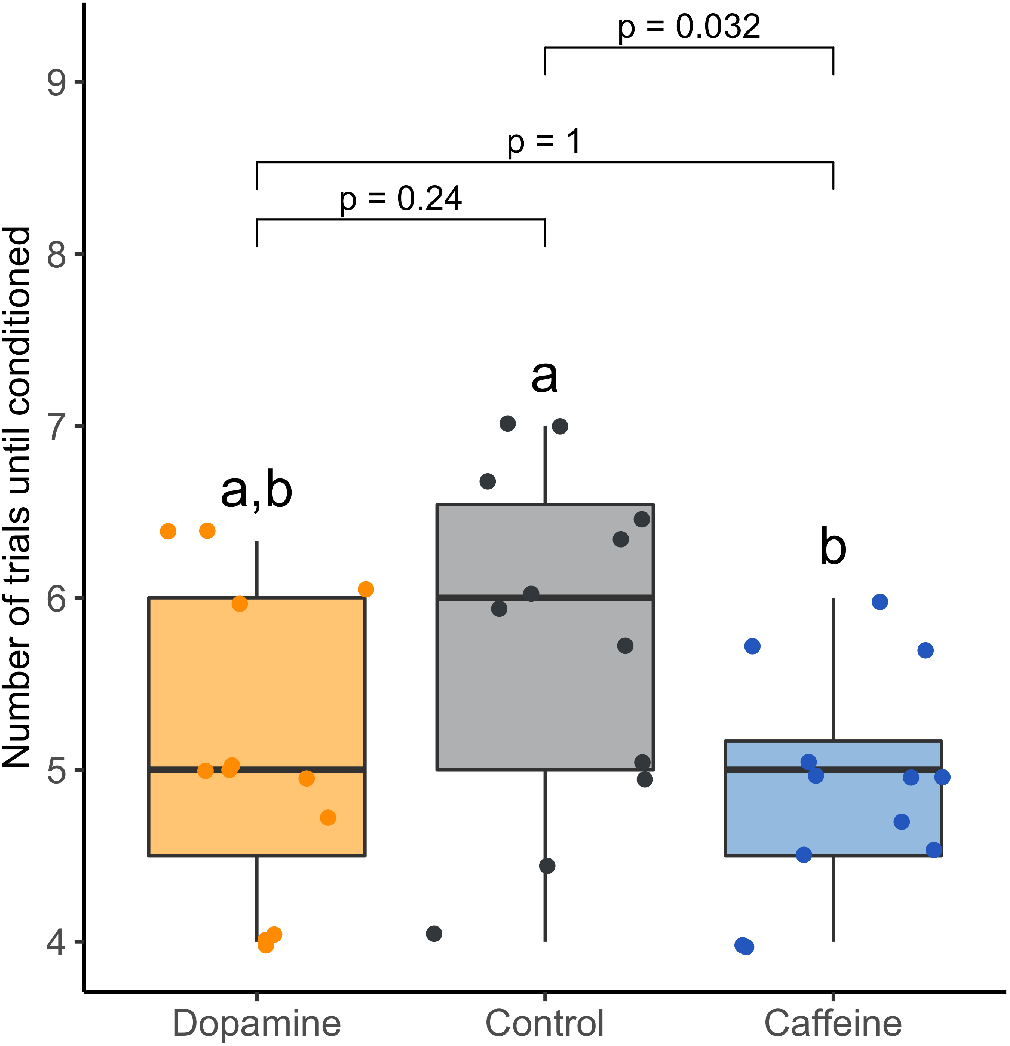
The effect of caffeine and dopamine on learning performance in the European honeybees, *Apis mellifera.* Caffeine-treated bees showed a higher learning performance than the control, requiring less training trials until conditioned to the stimulus. Significance was calculated using repeated measures anova and differences between groups were estimated using Multiple Comparisons of Means (Tukey Contrasts).

### 3.4 Classroom Discussion

Outreach projects are an essential tool to communicate and disseminate science to non-specific audiences. Although they usually lack a common structure, outreach activities are better internalised by the public when they are involved in major stages of data collection and discussion. As such, our practice gathered all major elements present on the scientific method: a) research subject was presented in a logical, sequential form using slides; b) based on background information provided, participants were instructed to build scientific hypothesis, study aims and predictions; c) data collection was performed based on a protocol and individuals were given full autonomy on their research; d) statistical analysis was conducted as group based on the results provided; e) all participants engaged on a collective discussion aiming to understand biological mechanisms that might have influenced the results; f) suggested new hypothesis and developments that could be implemented in a future event.

## 4. Discussion

### 4.1 Bee cognition

Caffeine is the most consumed psychoactive in the world, being used by different cultures and social groups to promote wakefulness. Similar to other psychoactive drugs, caffeine also affects dopamine signaling, by blocking dopamine transporters, stimulating its release from terminals and reducing reuptake [40–42]. Dopamine is a neurotransmitter with multiple functions, such as the control of reward-motivated behavior. Therefore, dopamine is an essential component in conditioning, memory and learning [43–45], being present in most multicellular animals [46]. Using the group-generated data, we were able to identify caffeine-induced improvements in honeybee learning when compared to dopamine alone. This was potentially due to caffeine increasing the dopamine production and reducing reuptake [40–42]. As a result, caffeine treated bees potentially had more available dopamine than dopamine treated bees.

It is important to note that this engagement protocol sacrifices data robustness for safety and reproducibility. For example, newly emerged bees have a greatly diminished aptitude for conditioning relative to older bees [1], but, as stated, lack venom and thereby are safer for classroom use. In addition, we used honeybees from a single hive. This pseudoreplication reduces the cost considerably and facilitates experimental setup as a single honeybee frame is used. In addition, concentrations of caffeine and dopamine much higher than those found naturally were employed [5,47–49]. This facilitated a drug-induced detectable phenotype. However, we acknowledge that these limitations and experimental design choices greatly reduce the generality and reliability of the data. Multiple groups of non-specialist participants can be leveraged to generate large datasets but these datasets should be viewed more as pilot studies. Any interesting findings should be repeated with a more granular experimental design.

### 4.2 Insect-based engagement

We propose using honeybees to highlight the importance of insects to global ecology and economic prosperity. Here we describe a straightforward and engaging activity that can be widely deployed to facilitate this. We do, however, appreciate that the sourcing and handling of bees is a potential issue in conducting this experiment. A potential solution to this is partnership with a local aparist or an apiarist society.

Insects are an essential part of most, if not all, ecosystems [50–53]. They provide multiple ecological functions, ranging from breaking down organic matter in the soil to pollination and the control of insect and plant pests [54–56]. Their diversity and abundance is directly related to the state of conservation of the environment, with more natural and undisturbed areas having a higher diversity and abundance than disturbed areas [57–60]. However, insects are now facing an unprecedented threat [61]. Worldwide their numbers have plummeted, but because of their size, and relative unimportance to the average citizen, scientists do not know the exact extent of their decline [61]. Educating the public by demystifying their presence, function and importance is imperative to solve this crisis [50,62,63]. Projects that stimulate contact and promote mutual respect between humans and invertebrates, such as citizen science, are more important today than ever [50,62,63].

Amongst insects, the European *Apis mellifera* is an ideal candidate for citizen science studies. They are likable, relatively docile (when carefully handled) and, because of their biology, they can have direct parallels with humans. For instance, they live in a society, they share food, communicate locations and even the necessity of grooming through the grooming dance [64]. Furthermore, their relatively larger brain, compared to another laboratory staple, the fruit fly *Drosophila melanogaster*, makes learning experiments simpler and easier to conduct with non-specialists [15,19].

### 4.3 Data Quality and Loss

The 20 percent loss in data observed in our experiment was well within what we consider acceptable given the complex experimental system employed. This loss is compensated by the increase in data points a project like this achieves. The experiment was conducted on a relatively small scale and thus, a limited amount of samples per experimental group was provided (3 bees per experimental group). Given the simplicity and low cost related to the experimental setup, a larger number of replicates could have been provided to the Ryukyu Girls group to compensate for any data loss.

### 4.4 Absence of adverse events

No adverse events, in particular bee stings, occurred during this activity. Honeybees pose an additional risk when compared to other model systems but when effectively managed this risk is greatly minimised. However, it is paramount that any honeybee-based engagement activities are mindful of the risks posed and take steps to mitigate these risks. Although strong advocates for the inclusion of honeybees in activities involving individuals untrained in their handling, we also recognise the need for restraining the bees.

### 4.5 Extension to other teaching scenarios

We propose that these methods could also be employed in classroom-based activities. With limited modification, our protocol could be extended to include teaching on hypothesis development and the scientific method generally. Moreover, lessons on data analysis, specifically in R, could also be included using markdown code we have shared. Finally, the additional training undergraduate students undertake throughout their education may also facilitate the more robust experimental designs discussed above without compromising safety.

## 5. Conclusion

With the ongoing worldwide concerns regarding Colony Collapse Disorder [65], we hope the use of *A. mellifera* in outreach, citizen science and education raises awareness and is instrumental in communities adopting more bee-friendly policies. We also hope that through engagement with sections of the communities typically absent from scientific discourse, we amplify this awareness and also foster lifelong critical learning.

## Funding

JAD, IK and MV were supported by the Okinawa Institute of Science & Technology. Experimental materials were supported by the Okinawa Institute of Science & Technology and MV. Subsequent analysis and manuscript collation was supported by JSPS KAKENHI Grants 19K06795 and 19K16205 awarded to JAD and MV respectively. The Ryukyu Girls program and participants were supported by Okinawa Institute of Science & Technology Graduate University, University of the Ryukyus, Okinawan Prefectural Board of Education, Okinawan Prefectural Government, the Okinawa Institute of Science and Technology Graduate University Promotion Council and the ‘FY2018 GST Female Jr. high and high school student support program..

## Author Contributions

MV designed and developed the honeybee experimental framework with input from all authors. JAD, IK, MT & MV conducted the engagement activity. All authors wrote and edited the paper.

## Acknowledgment

We would like to acknowledge the support of the Okinawa Institute of Science and Technology Diversity Section. We would also like to thank all participants of the 2018 Ryukyu Girls program for their enthusiasm and dedication.

## Notes

https://github.com/marivelasque/HoneybeeOutreach.git

### Competing Interest Statement

The authors have declared no competing interest.

